# Imaging live bacteria at the nanoscale: comparison of immobilisation strategies

**DOI:** 10.1101/685024

**Authors:** Georgina Benn, Alice L. B. Pyne, Maxim G. Ryadnov, Bart W Hoogenboom

## Abstract

Atomic force microscopy (AFM) provides an effective, label-free technique enabling the imaging of live bacteria under physiological conditions with nanometre precision. However, AFM is a surface scanning technique, and the accuracy of its performance requires the effective and reliable immobilisation of bacterial cells onto substrates. Here, we compare the effectiveness of various chemical approaches to facilitate the immobilisation of *Escherichia coli* onto glass cover slips in terms of bacterial adsorption, viability and compatibility with correlative imaging by fluorescence microscopy. We assess surface functionalisation using gelatin, poly-L-lysine, Cell-Tak™, and Vectabond^®^. We describe how bacterial immobilisation, viability and suitability for AFM experiments depend on bacterial strain, buffer conditions and surface functionalisation. We demonstrate the use of such immobilisation by AFM images that resolve the porin lattice on the bacterial surface; local degradation of the bacterial cell envelope by an antimicrobial peptide (Cecropin B); and the formation of membrane attack complexes on the bacterial membrane.

## Introduction

Live single-cell imaging can advance the current understanding of cellular heterogeneity in bacterial populations at the level of an individual cell as a function of time. Although the results of traditional cell culture measurements dealing with large cell numbers are statistically significant, they cannot address the behaviour of individual cells because of the averaging on which they rely. Higher-resolution methodologies are necessary to access single-cell analysis and complement these measurements^1^. High-resolution imaging techniques with integrated microfluidic devices and cell tracking software have provided qualitatively new insights into cellular processes. For example, fluorescence microscopy used for single-molecule tracking inside live bacteria helped to reveal that the *lac* transcription factor finds its binding site via the facilitated diffusion model^2^. Microfluidic devices are also powerful tools for single cell bacterial analysis. For example, microfluidic devices have been combined with cell tracking, to enable the rapid detection of antibiotic resistance in clinical isolates in under 10 minutes^3^. Arguably, however, atomic force microscopy (AFM) is the technique of choice for accurate and label-free molecular cell measurements^4–9^.

Indeed, AFM performed in water or physiological buffers, including cell culture media, allows the acquisition of nanometre resolution images with no ensemble averaging, under physiological conditions^10^. The principle of AFM is to use a sharp tip on a cantilever to directly probe the features of an analyte immobilised on a surface. By scanning across the surface of the analyte the cantilever allows the build-up of a contour or topography map of the surface features, line by line (Figure 1). Notable examples of AFM for cell imaging revealed the dynamics of filopodia on live hippocampal neurons^6^; a net-like structure of porins in the outer membranes of Gram-negative bacteria^4,11^; and differences in action of antimicrobial peptides on bacteria in water or LB broth^12^ and mechanistic insights into new poration mechanisms^13^. However, regardless of application, AFM, like other single-cell techniques, relies on cells remaining immobilised to a substrate.

**Fig. 1.**
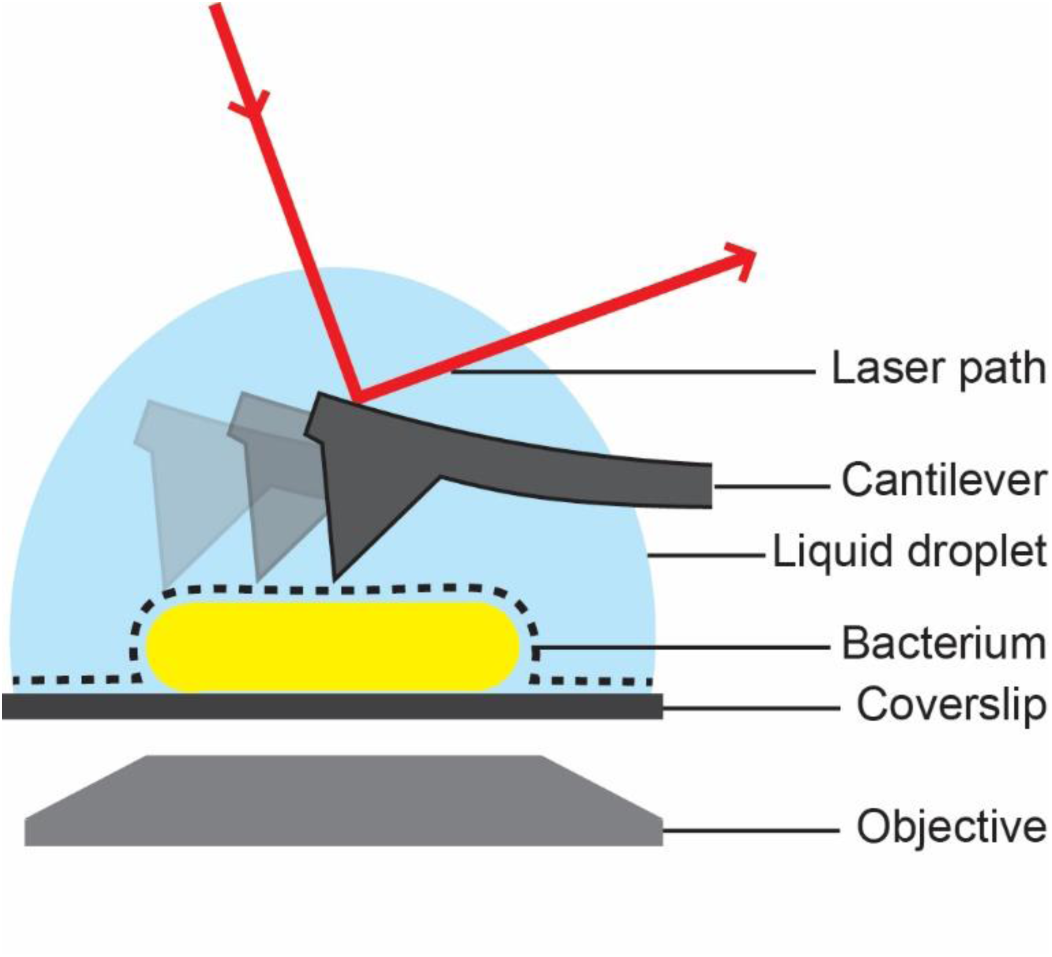
Schematic of bacterial cell attached to a glass substrate for microscopic analysis in solution. Inverted optic microscopy and complementary fluorescence microscopy (via objective below) can be used to find and inspect cells at low resolution. For AFM analysis, the bacterial surface is traced by a sharp needle on a flexible cantilever. The bending of the cantilever is a measure of the force between the surface and the AFM probe, this is detected via the deflection of a laser beam.

The physical nature of AFM means that cells must be stably adhered to a substrate. Others have used different methods to promote bacterial adhesion for AFM imaging with variable successes. Firstly, microfluidics can be used to physically trap bacteria in wells^14^. This can be achieved in a range of buffers including growth media and requires no chemical interactions between substrate and bacteria, thus leaving cell viability unaffected. However, the trapping of cells requires appropriately sized holes, which itself depends on the species of bacteria. *E. coli* and *B. subtilis* have been immobilised using a microfluidic device that also allowed simultaneous fluorescence imaging, but the fabrication of these devices is time consuming^12,14^. To immobilise *Mycobacterium* species, polycarbonate filters can be used^15,16^, but the efficiency of this approach was reported as low and was not feasible for most species due to their size^17^. A covalent attachment of bacteria to a surface could affect cell viability and should be avoided^14^. Generally, mica is the most common substrate for AFM as it can easily be cleaved to provide an atomically flat surface^18^ For cell imaging, however, it is more convenient to have visual pre-scanning by an inverted optical microscope, so glass coverslips or slides are used to find and select cells for high-resolution AFM imaging^19,22^. Ensuring the adherence of cells onto glass or mica is a prerequisite for AFM sample preparation. An ultimately reliable immobilisation method would be compatible with physiological buffers and have no effects on cell viability or morphology. Furthermore, such a method should meet time considerations of AFM imaging, particularly when visualising cellular or cell-related processes over prolonged periods of time. Thus, the choice of immobilisation methods for accurate and reliable AFM imaging is limited to those that can satisfy the fairly stringent suitability requirements for sample preparation.

Here we compare four adhesion methods for two different strains of *E. coli*. This bacterium is one of the most common Gram-negative pathogens. We focus on Gram-negative cells because they are clinically important-being responsible for a significant burden to healthcare worldwide ^20^ and have been used extensively in AFM studies of bacteria^4,11,12,21,22^ so far.

## Materials and methods

### Bacterial strains and preparation

For mid-log phase bacteria, an *E. coli* MG1655 (provided by the Rooijakkers lab, University Medical Centre Utrecht) or BL21 colony was picked from a LB-agar plate and grown in 3 mL LB broth (Lennox) for 3 hours at 37°C in a shaking incubator. 500 μL of culture was then spun at 5000 rpm for 2 minutes, the supernatant removed and bacteria resuspended in 500 μL of HEPES buffer (20 mM HEPES, 120 mM NaCl, pH 7.4), PBS (10 mM phosphate buffer, 137 mM NaCl, 2.7 mM KCl, pH 7.4), PB (10 mM phosphate buffer, pH 7.4) or millliQ water (mQ). Spinning and resuspension was repeated 3 more times to remove all LB.

### Glass cleaning

13 mm glass coverslips (VWR) were placed in a rack and rinsed in a stream of mQ. They were then sonicated in 2% SDS at 37 kHz and 100% power in a Fisherbrand™ bath sonicator (Fisher Scientific) for 10 minutes. Next, they were rinsed and soaked in mQ, followed by ethanol and dried with nitrogen. They were then plasma cleaned at 70% power for 2 minutes in a plasma cleaner (Zepto, Diener Electronic). The whole procedure was then repeated once more and coverslips functionalised as described below. Coverslips were used immediately after preparation and not stored.

### Bacteria immobilisation

100 μL of bacteria in HEPES, PBS, PB or mQ was added to each fully prepared coverslip (see below) and incubated at room temperature for 15 minutes on gelatin, 5 minutes on PLL and 30 minutes on Cell-Tak™ or Vectabond^®^. Unadhered bacteria were washed 3 times by rinsing in 1 mL of an appropriate buffer. Care was taken to avoid drying the sample out at any point. It is worth noting that Vectabond^®^ coated glass is hydrophobic: extra care was taken not to dislodge the droplet.

### Glass functionalisation

#### Gelatin

Gelatin solution was prepared by adding 0.5 g of gelatin (G6144, Sigma) to 100 mL of mQ water just off the boil. The mixture was then swirled until all gelatin had dissolved and the temperature had dropped to 60-70°C^23^. Freshly cleaned coverslips were then dipped into the warm gelatin, removed and balanced on their edges until dry. Coverslips were then glued to clean glass slides using biocompatible glue (Reprorubber thin pour, Flexbar, NY). Bacteria were added as described above.

#### Poly-L-Lysine

Clean glass coverslips were placed flat on a clean slide and a 100 μL droplet of 0.01% poly-L-lysine (P4832, Sigma) was added. After 5 minutes at room temperature, the coverslips were rinsed in a stream of mQ, dried in nitrogen and glued to clean glass slides using biocompatible glue. Bacteria were added as described above.

#### Cell-Tak™

Clean coverslips were glued to glass slides using biocompatible glue. A Cell-Tak™ solution was then prepared by mixing 1 mL 0.1 M sodium bicarbonate, pH 8.0 with 40 μL Cell-Tak™ (BD Diagnostics, USA) and 20 μL 1M NaOH. 100 μL of this solution was applied to a glued down coverslip and incubated for 30 minutes at room temperature. Coverslips were then rinsed with a stream of mQ and nitrogen dried. Bacteria were added as described above.

#### Vectabond^®^

Cleaned coverslips were put into a rack and submerged in 50 mL acetone for 5 minutes, then moved to a 50:1 solution of Acetone:Vectabond^®^ (Vector Laboratories, USA) for 5 minutes. Finally, coverslips were dipped several times in mQ, nitrogen dried and glued to clean glass slides using biocompatible glue. Bacteria were added as described above.

### Determining bacterial adhesion and survival

Bacteria were imaged immediately after immobilisation (data not shown) and two hours after immobilisation, using an Andor Zyla 5.5 USB3 fluorescence camera on an Olympus IX 73 inverted optical microscope. Cell death of bacteria was assessed by adding 1 μL of SYTOX™ green nucleic acid stain (S7020, Sigma) to the sample to mark dead cells. Brightfield and fluorescence images were taken of the same region to calculate the number of cells adhered and the percentage of those that were dead. Images used in Figures 2 and 3 have been cropped and the contrast enhanced in FIJI-ImageJ^24^ to show bacterial cells more clearly.

**Fig. 2.**
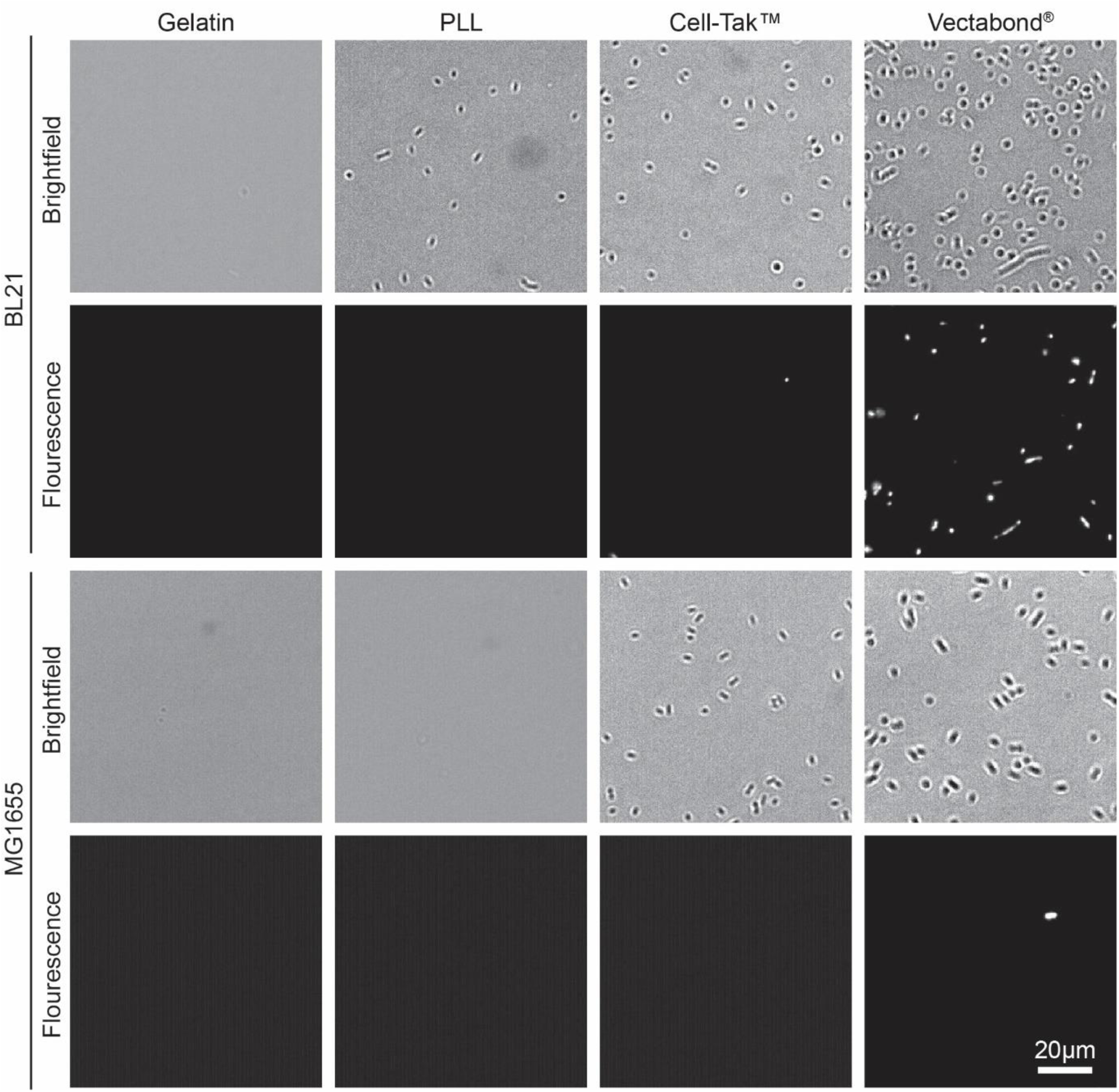
Representative brightfield and fluorescence images of E. coli cells (BL21 and MG1655) immobilised on different coatings in HEPES buffer. Fluorescent bacteria are labelled with SYTOX^®^ green dead cell stain.

**Fig. 3.**
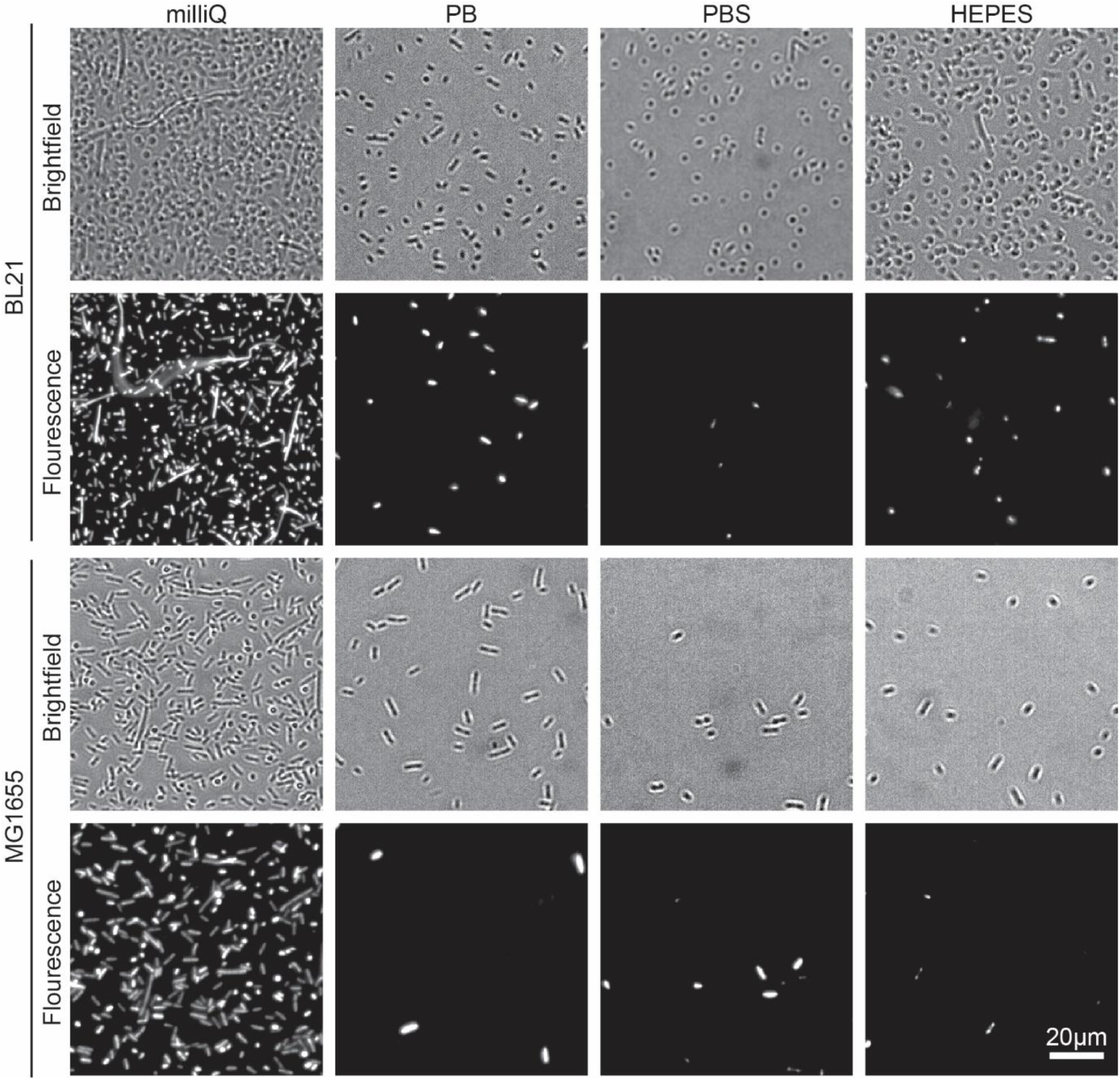
Representative brightfield and fluorescence microscopy images of E. coli cells (BL21 and MG1655) immobilised on Vectabond^®^ coated coverslips under different buffer conditions. Fluorescent bacteria are labelled with SYTOX^®^ green dead cell stain.

Image analysis was performed using FIJI-ImageJ^24^, with settings and parameters as follows. The number of bacteria in brightfield and SYTOX™ images was calculated by cropping each image, then picking bacteria using ImageJ macros. Depending on the quality of image and number of bacteria in each field of view, bacteria were picked using different procedures. The effectiveness of image processing was assessed by comparison with original images. Generally, brightfield images were smoothed, converted to binary and despeckled to remove noise. To remove large background features, bacteria were identified using the ‘find edges’ function, or a background subtraction (rolling ball radius of 25 pixels, pixel size 0.32 µm/pixel) was applied. For SYTOX™ images, a threshold was applied (either by the Otsu or Default method) and the image despeckled. When the density of bacteria was high, a watershed algorithm was used to identify individual cells; and when there were no bacteria in the cropped image, it was not processed. The number of cells was counted as the number of particles with an area between 2 and 300 pixels, corresponding to approximately 0.6 to 100 µm^2^. The field of view was 360 × 240 µm^2^. The number of cells counted was plotted using Origin (OriginLab, MA, USA).

### Peptides and proteins

After immobilising bacteria, sample surfaces were blocked by incubation with 20 mM HEPES, 120 mM NaCl, 2.5 mM MgCl_2_, 0.1% BSA (HEPES/BSA) for 30 minutes at room temperature, the samples were then washed with 1 ml HEPES buffer three times. An antimicrobial peptide cecropin B was added to bacteria to a final concentration of 5 µM^25^. To image the membrane attack complex on bacteria, components of the MAC were added sequentially as described elsewhere^4^. Briefly, a 10% solution of C5 deficient serum (CompTech, Texas USA) in HEPES/BSA was added to bacteria and incubated for 20 minutes at 37°C, the sample was then washed to remove serum. 0.1 µg/mL of each MAC component in HEPES/BSA were then added in two stages: C5, C6 and C7 (provided by the Rooijakkers lab, University Medical Centre Utrecht) were added, incubated for 5 minutes and washed; then C8 (CompTech, Texas USA) and C9 (provided by the Rooijakkers lab, University Medical Centre Utrecht) were added for 20 minutes and washed. The samples were imaged by AFM in tapping mode as described below.

### Atomic force microscopy

AFM was performed in intermittent-contact mode on a Nanowizard III AFM with an UltraSpeed head (JPK, Germany; now Bruker AXS, CA, USA) using a FastScanD (Bruker AXS, CA, USA) cantilever with 0.25 N/m spring constant and 120 kHz resonance frequency. All AFM was performed in liquid in HEPES buffer and was performed within 3 hours of immobilising. Images are 512×512 pixels (unless otherwise specified). 5×5 µm^2^ scans were performed at a line frequency of 1 Hz, 500×500 nm^2^ and 350×350 nm^2^ scans were performed at 3-5 Hz. Data was analysed in Gwyddion 2.52 (http://gwyddion.net/)^26^. 5× 5 µm^2^ scans were processed by applying a first-order plane fit. A first-order plane fit, followed by line-by-line 2^nd^ order flattening and a gaussian filter with σ = 1 pixel, to remove high-frequency noise, was applied to 500×500 nm^2^ and 350× 350 nm^2^ scans.

## Results and Discussion

In this study, we have looked at two strains of *E. coli* (BL21 and MG1655) in four different buffers (milliQ water, PB, PBS and HEPES), for four different functionalisation techniques (Gelatine, PLL, Cell-Tak™ and Vectabond^®^). The first stage of coverslip preparation was an extensive cleaning process that is essential in achieving high immobilisation efficiency. *E. coli* BL21 and MG1655 were used because they are common model strains for single cell studies^3,4,8,12,21,27,28^. They also differ in their outer membrane structures; while MG1655 has a higher abundance of flagella and the presence of the polysaccharide region of the lipopolysaccharide (LPS), BL21 has fewer flagellar and no LPS^29,30^.

The efficiency of bacteria adhesion on selected coatings was quantified by counting the total number of bacteria per unit area (360 × 240 µm^2^) using brightfield microscopy. Cell viability was verified by the fluorescence of the nucleic-acid dye SYTOX™, where fluorescence is a signature of permeability of the cell envelope and bacterial death.

The efficiency of bacterial adhesion onto glass was highly variable. This variability was observed between different strains of *E. coli*, between different surfaces (Figure 2 demonstrates the degree of variation between surface types in HEPES) and between different buffers (Figure 3 demonstrates the degree of variation between buffers on Vectabond^®^). These variations are quantified for all conditions in Figure 4a.

**Fig. 4.**
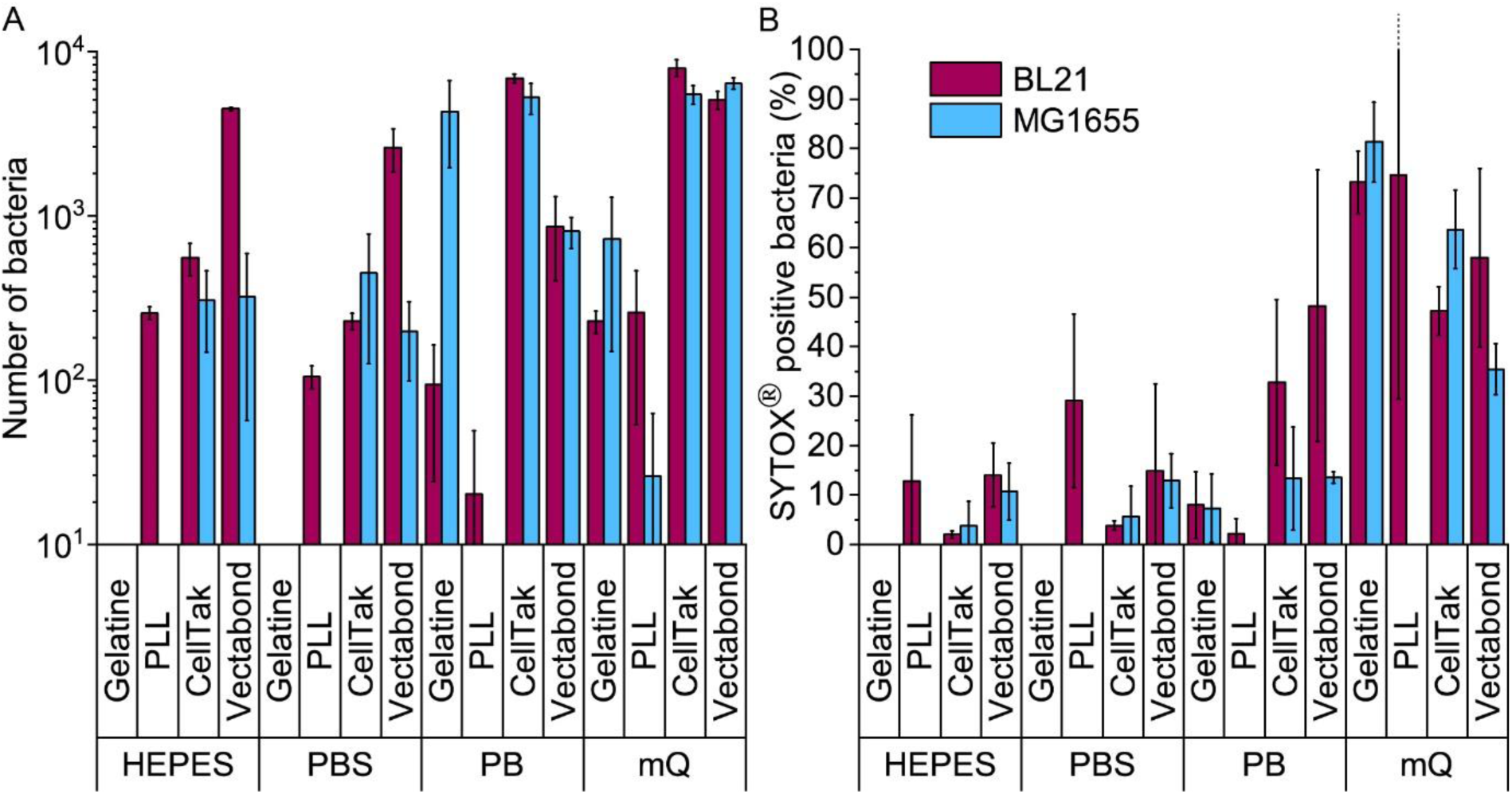
(A) The mean number of all bacteria (live and dead) in a 360 × 240 µm^2^ field of view for each condition tested. Note the logarithmic scale on the vertical axis. (B) The mean percentage death of bacteria in a field of view for each condition tested, dead bacteria were identified as SYTOX^®^ positive cells. Error bars are standard deviations of the mean (n=3).

We found that MG1655 *E. coli* are more difficult to immobilise than BL21. In Figure 4a, we see that the number of BL21 bacteria adhered was greater than that of MG1655 in all but 4 of the 16 conditions tested. This is possibly due to the fact that BL21 lack the LPS^29^; or due to a higher abundance of flagellar on MG1655, which increases the motility of this strain^30^.

The buffer composition also affects the immobilisation of bacteria. Figure 4a shows that, compared to milliQ water, bacteria are less adherent in low salt buffer (PB) and even less adherent in high salt buffers (PBS and HEPES). The exception to this is BL21 *E. coli* on PLL, where the adhesion is lower in PB than PBS or HEPES. In milliQ, adhesion is high for both strains on all surfaces (>100 bacteria per image 360 × 240 μm^2^), except MG1655 on PLL which leads to ∼30 bacteria per image.

Different immobilisation techniques also yield different levels of adhesion and are affected by buffer composition to different degrees (Figure 4a). Gelatin and PLL are the most common methods used for immobilisation of bacteria^17,31^. These are cationic protein coatings that promote the attachment of anionic bacteria via electrostatic interactions. The effectiveness of these coatings was found to be highly dependent on buffer conditions and *E. coli* strain. For gelatin, no bacteria were adhered to coverslips unless bacteria were immobilised in milliQ or PB, when adhesion is high. This may be due to the masking of electrostatic interactions by monovalent ions^17^. Furthermore, the preparation of gelatin coated coverslips is time consuming, since air drying of coverslips takes hours and may complicate the planning and design of AFM experiments, in particular those that require prolonged scanning. For these reasons, we do not recommend the use of gelatin to adhere bacteria.

In contrast, PLL requires the shortest preparation time and is a relatively cost-effective option. In the case of MG1655 on PLL, adhesion was poor in all conditions including milliQ possibly due to the flagella^32^ and polysaccharides on the surfaces of these cells^29^. However, adhesion of BL21 onto PLL was good (>100 bacteria per image) in high salt buffers and milliQ, but poor in PB. This is contrary to the poor adhesion of bacteria in phosphate buffers on gelatin and may be because PLL has larger net positive charge than gelatin^31^.

The third immobilisation technique used was Cell-Tak™. Cell-Tak™ is an acidic solution of polyphenolic proteins purified from marine mussels. When neutralised with sodium bicarbonate, the proteins absorb onto a surface, coating a glass coverslip for bacteria to adhere to^33^. Brightfield images demonstrated good adhesion of both strains of bacteria in all conditions, supporting previous work showing good adhesion for a range of bacteria, even in nutrient broth^17^. On Cell-Tak™ adhesion was high for both strains of bacteria in all conditions (Figure 4a).

Finally, Vectabond^®^ is a solution predominantly made up of 3-Aminopropyltriethoxysilane (APTES)^34^ which coats coverslips with amine groups and is believed to adhere bacteria via electrostatic and hydrophobic interactions^17,35^. This is similar to gelatin and PLL coatings. However, adhesion of bacteria onto Vectabond^®^-coated coverslips gave greater numbers of adhered cells in all conditions, in particular, the coating supported high levels of MG1655 adhesion in buffers.

Figure 4b shows the percentage of dead bacteria in each image. As with adhesion, cell death depends on the bacterial strain: the proportion of dead BL21 was lower in only 3 of the conditions tested, compared to the survival of MG1655 in the same conditions. Buffer composition also affects cell survival: immobilisation in milliQ consistently led to a high percentage of dead cells (35-82% dead). Bacteria in HEPES and PBS survived better than cells in PB when immobilised on Cell-Tak™ and Vectabond^®^, but not on PLL where survival was approximately equal in PB, PBS and HEPES. For gelatin and PLL, when bacteria were adhered, the percentage cell death was low in all buffers except milliQ.

Next, we carried out AFM on bacteria immobilised in HEPES (Figure 5), HEPES was used because survival was good for BL21 and MG1655 on all surfaces (Figure 4b). BL21 on PLL were well adhered, bacteria were smooth and resolution was high enough to see porins (∼7 nm pores in the outer membrane^4^) covering the surface. When imaging bacteria adhered with Cell-Tak™, unidentified aggregates approximately 10-20 nm high and 50-100 nm wide were observed on both BL21 and MG1655, making the samples unusable for high resolution studies. We note that Cell-Tak™ has previously been used in previous studies^17^ and in our own published^4^ and unpublished research, without this problem of aggregates. Hence, it cannot be ruled out as a viable immobilisation strategy – here we just report the risk of aggregation issues. This problem also highlights the importance of AFM based experiments, since cells appear unchanged when looking at brightfield images and there is no increase in cell death. Finally, MG1655 bacteria on Vectabond^®^ coated coverslips were well adhered and usable for high resolution imaging.

**Fig. 5.**
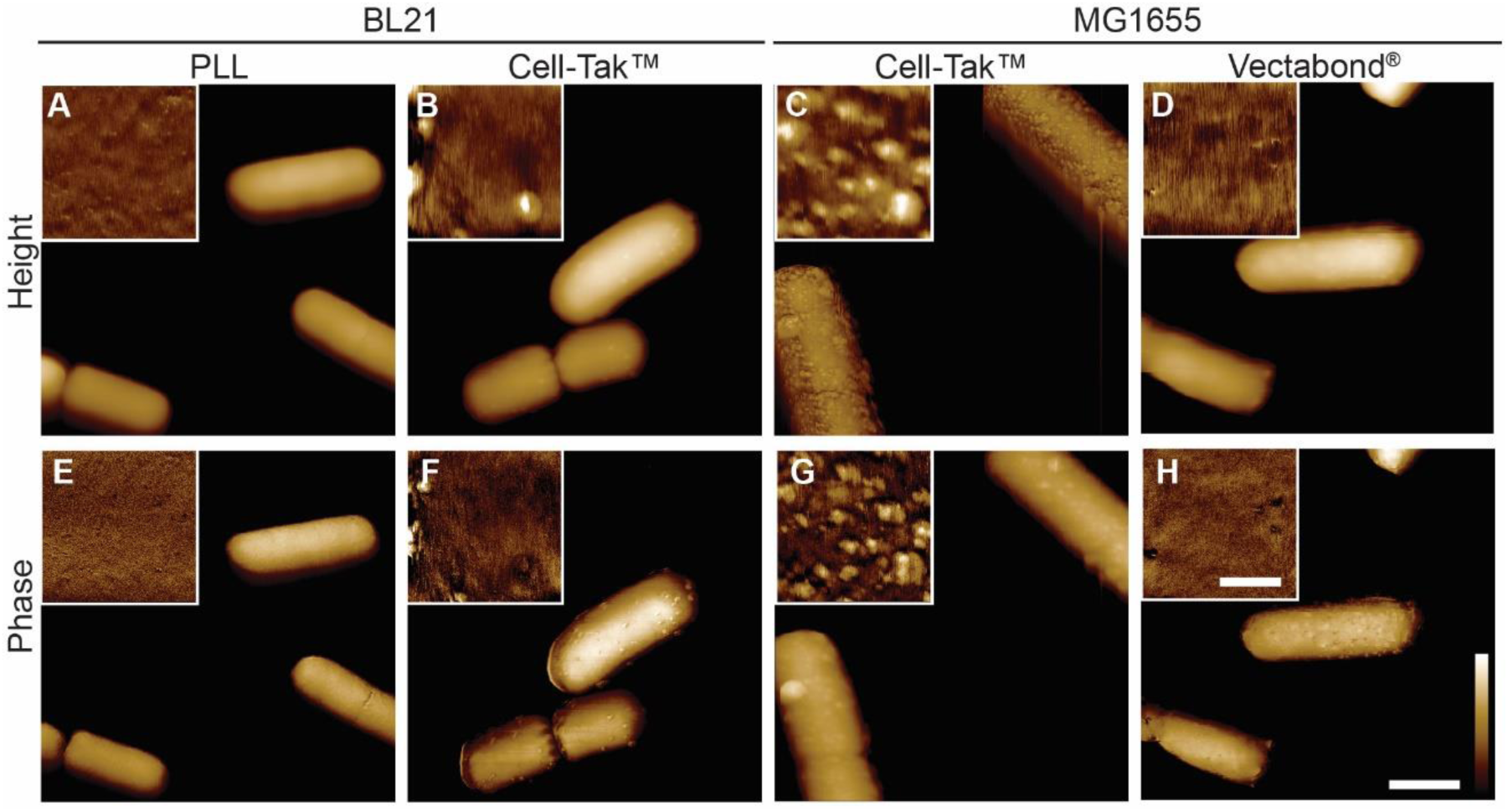
Tapping mode atomic force microscopy height and phase images of E. coli bacteria immobilised onto glass coverslips. Lateral scale bar (A-H) 1 µm, insets 250 nm. Vertical colour scales (A) 600 nm, inset 10 nm; (B-C) 600 nm, insets 30 nm; (D) 600 nm, inset 10 nm; (E) 10 deg, inset 1.5 deg; (B-C) 10 deg, inset 2 deg; (D) 10 deg, inset 1.5 deg.

To demonstrate the performance of AFM on the adhered bacteria, Figure 6 shows 350×350 nm^2^ scans of the surface of *E. coli* bacteria. Figure 6A shows a pattern of ∼10 nm wide pits at the surface of MG1655 *E. coli*, similar to previous observations^4,11^. The dimensions of this pattern are consistent with those observed for porins on isolated outer membranes^36^. This indicates we resolve the outer membrane porin lattice on live *E. coli*. Figure 6B shows the degradation of the *E. coli* surface due to an antimicrobial peptide that is known to target both Gram-positive and Gram-negative bacteria, cecropin B (CecB)^25^. Finally, Figure 6C shows pores that have been assembled following exposure of *E. coli* to the immune proteins that form bactericidal membrane attack complexes (MACs). The size and shape of these rings is consistent with cryoEM and AFM data of the whole MAC pore inserted into lipid bilayers^37,38^.

**Fig. 6.**
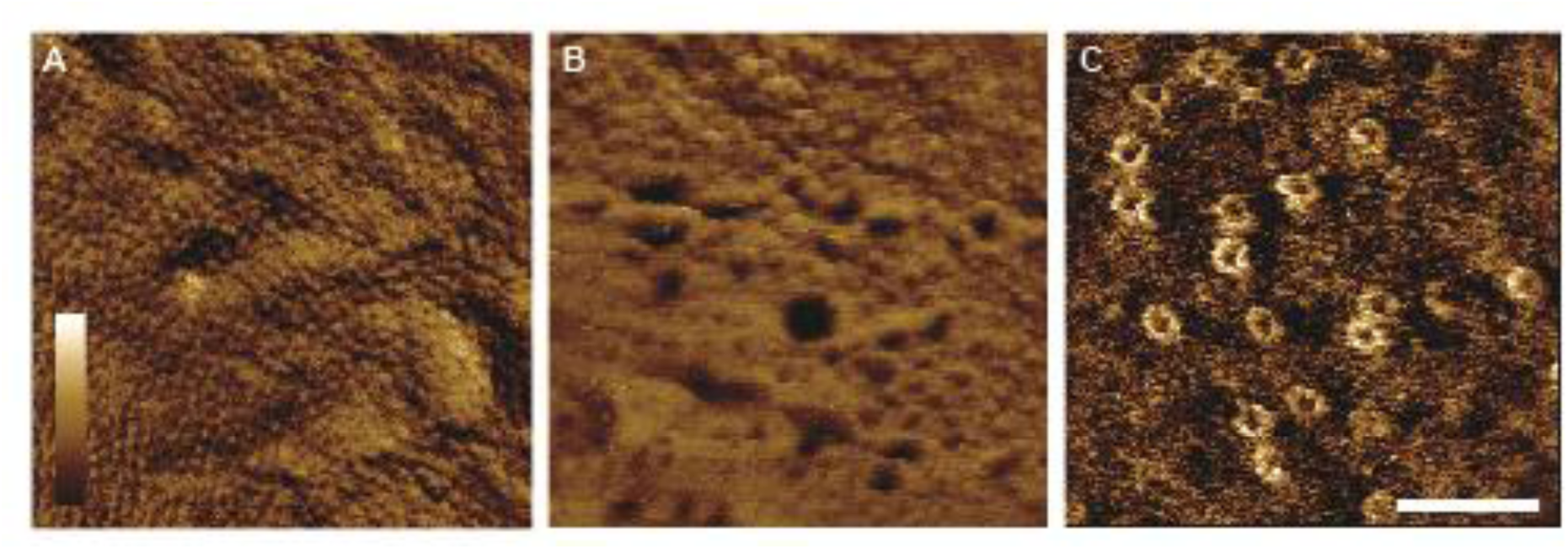
Tapping mode atomic force microscopy phase images of MG1655 (A) and BL21 (B-C) E. coli bacteria immobilised onto glass coverslips. (A) When bacteria, on Vectabond^®^ coated coverslips in HEPES buffer, are imaged at high resolution, a network of porins can be seen in the outer membrane. (B) AFM can be used to investigate the mechanism of action of antimicrobial peptides. As an example, 5 µM Cecropin B was applied to bacteria immobilised onto Vectabond^®^ coated coverslips in HEPES buffer, resulting in nanometre-scale poration of the outer membrane. (C) Using bacteria here immobilised on PLL in HEPES buffer, the formation of the membrane attack complex (MAC) can be investigated on live bacteria. The MAC pores can be observed as rings in the membrane. Lateral scale bar is 100nm. Vertical colour scale is (A) 2 deg (B) 4 deg and (C) 3 deg. (A-B) are 512×512 pixels, (C) is 256×256 pixels.

## Conclusions

The development of a robust immobilisation technique is an essential part of any AFM experiment. As well as efficiency, important considerations include time, cost and reliability. Another consideration is the impact of the surface functionalisation on the following AFM experiments. We have found that buffers are essential to keep bacteria viable for prolonged time periods however, they tend to reduce the efficiency of immobilisation. Successful immobilisation methods were Vectabond^®^ for all conditions tested and PLL in some conditions. By contrast, gelatin was the least successful immobilisation technique in all buffered conditions tested. We also highlight the importance of performing AFM on bacteria before deciding on an immobilisation technique, since Cell-Tak™ can appear successful until AFM is performed and bacteria may be coated with an unknown aggregate. Finally, we show some examples of images obtained by AFM that show high-resolution, *in situ* changes to the surface of live bacteria.

## Conflicts of interest

There are no conflicts to declare

## Acknowledgements

We thank Edward Parsons and Isabel Bennett for their advice on immobilisation protocols; Kate Hammond (NPL) for Cecropin B; Dani Heesterbeek and Suzan Rooijakkers (Utrecht) for providing MAC proteins and advice; and the UK Engineering and Physical Sciences Research Council (EP/N509577/1) and the UK Medical Research Council (MR/R000328/1 and MR/R024871/1) for funding.

